# Modeling the consequences of regional heterogeneity in human papillomavirus (HPV) vaccination uptake on transmission in Switzerland

**DOI:** 10.1101/125518

**Authors:** Maurane Riesen, Victor Garcia, Nicola Low, Christian L. Althaus

## Abstract

**Background:** Completed human papillomavirus (HPV) vaccination by age 16 years among women in Switzerland ranges from 17 to 75% across 26 cantons. The consequences of regional heterogeneity in vaccination coverage on transmission and prevalence of HPV-16 are unclear.

**Methods:** We developed a deterministic, population-based model that describes HPV-16 transmission among young adults within and between the 26 cantons of Switzerland. We parameterized the model using sexual behavior data from Switzerland and data from the Swiss National Vaccination Coverage Survey. First, we investigated the general consequences of heterogeneity in vaccination uptake between two sub-populations. We then compared the predicted prevalence of HPV-16 after the introduction of heterogeneous HPV vaccination uptake in all of Switzerland with homogeneous vaccination at an uptake that is identical to the national average (52%).

**Results:** HPV-16 prevalence in women is 3.34% when vaccination is introduced and begins to diverge across cantons, ranging from 0.14 to 1.09% after 15 years of vaccination. After the same time period, overall prevalence of HPV-16 in Switzerland is only marginally higher (0.55 %) with heterogeneous vaccination uptake than with homogeneous uptake (0.49%). Assuming inter-cantonal sexual mixing, cantons with low vaccination uptake benefit from a reduction in prevalence at the expense of cantons with high vaccination uptake.

**Conclusions:** Regional variations in uptake diminish the overall effect of vaccination on HPV-16 prevalence in Switzerland, although the effect size is small. Cantonal efforts towards HPV-prevalence reduction by increasing vaccination uptake are impaired by cantons with low vaccination uptake. Harmonization of cantonal vaccination programs would reduce inter-cantonal differences in HPV-16 prevalence.

## 1. Introduction

The first vaccine against human papillomavirus (HPV) was licensed in 2006 and is now widely used in many countries. At the population-level, HPV vaccination has led to a substantial reduction in the prevalence of the targeted HPV types (HPV-16/18/6/11 for the quadrivalent vaccine) as well as anogenital warts [1]. Most vaccination programs target girls or young women before they become sexually active. Regional differences in vaccination uptake have emerged in some countries after implementation of the vaccination programs [2, 3]. These differences are very pronounced in Switzerland where the proportion of 16 year old girls completing the three dose vaccination schedule ranges from 17 to 75% in 26 cantons (states) (Fig. 1) [4, 5]. The cantonal heterogeneity in vaccination uptake can be partly explained by differences in the way the vaccine is offered to girls and young women (e.g., school-based programs, general practitioners or gynecologist). Other factors, such as cultural differences between the cantons might play a role too. To date, the potential epidemiological consequences of regional variation in vaccination uptake on transmission and prevalence of HPV in Switzerland and other countries are not well understood.

**Fig. 1.**
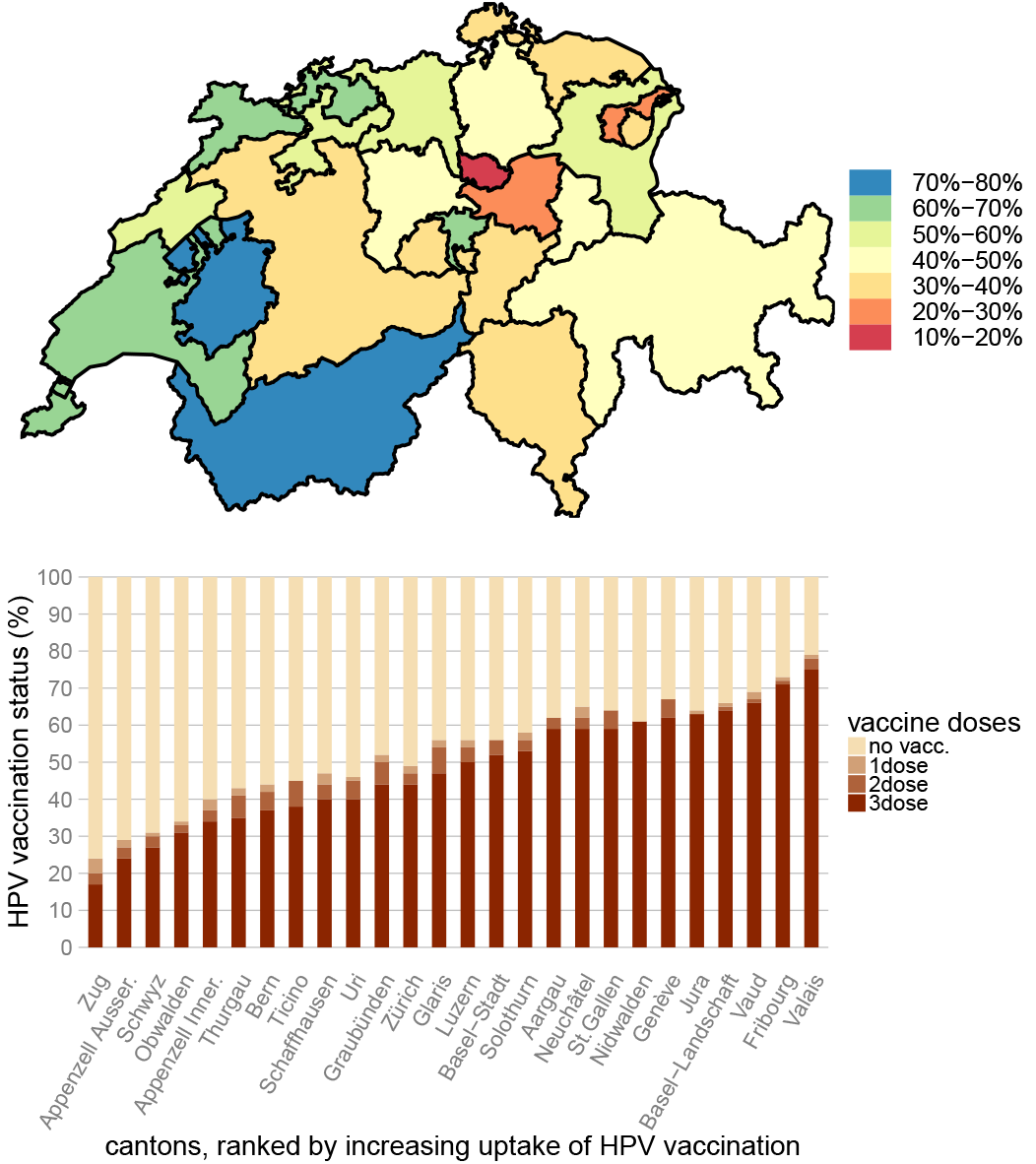
HPV vaccination uptake in 16 year old girls in Switzerland. Data represent the last completed survey period (2011–2013) of the Swiss National Vaccination Coverage Survey (SNVCS). Data for Geneva and Appenzell Innerrhoden are from 2010 and 2014, respectively.

Mathematical models have played an important role in estimating the expected impact of vaccination on the transmission of HPV [6–8] and other infections [9]. Investigating the consequences of spatial heterogeneity in vaccination uptake has received less attention, with some mentionable exceptions. Studies on measles vaccination [10, 11] and canine rabies [12] showed that spatial vaccination heterogeneity leads to less effective control of the targeted disease when compared with homogeneous vaccination. The debate about heterogeneity in HPV vaccination uptake has focused on sex-specific vaccination [13, 14]. Sex-specific vaccination is expected to be more beneficial than homogeneous (male/female) vaccination in a heterosexual population because both sexes are required for transmission. Therefore if only one sex is targeted by the vaccine, although around 50% of the total population would be vaccinated, the transmission would be blocked as the vaccine-targeted sex would act as a dead-end host. Spatial variation in HPV vaccination uptake between states in the United States of America (USA) has been taken into account in a modeling study that quantified the epidemiological impact and cost-effectiveness of adopting a new, nonavalent HPV vaccine [15]. This study illustrated that expanding vaccination coverage in states with low coverage would result in the greatest health impact because of the decreasing marginal returns of herd immunity. This finding is supported by another modeling study from Canada showing that the effect of unequal vaccination uptake among school girls by ethnicity on cervical cancer incidence may be lower than with equal vaccination [16]. The effects of spatial heterogeneity in vaccination uptake crucially depend on sexual mixing between different regions, as well as herd immunity thresholds and other disease-specific characteristics. A better understanding of how these factors affect the transmission and prevalence of HPV may help to better interpret the expected or observed impact of HPV vaccination programs.

The aim of this study was to investigate the impact of heterogeneous vaccination uptake and different sexual mixing scenarios on the prevalence of HPV-16 in Switzerland. We developed a mathematical model of HPV-16 transmission among young heterosexual adults. We parameterized the model using Swiss sexual behavior data and calculated the pre-vaccination prevalence and the basic reproduction number (*R*_0_) of HPV-16. First, we investigated the general consequences of heterogeneous vaccination uptake in a simple model with two sub-populations. We then simulated the transmission of HPV-16 within and between the 26 cantons of Switzerland assuming three different scenarios for inter-cantonal sexual mixing. We compared the predicted post-vaccination prevalence of HPV-16 after the introduction of heterogeneous HPV vaccination uptake with a default scenario of homogeneous vaccination.

## 2. Methods

### 2.1. HPV-16 transmission model

We developed a deterministic, population-based model of HPV transmission that is based on well-establish work on modeling sexually transmitted infections (STIs) [17–19]. For simplicity, we focused on HPV-16 only as it is the most common oncogenic type in women worldwide [20] and responsible for more than 50% of invasive cervical cancers [21]. We implemented the spatial (cantonal) structure into a meta-population model, and considered the population of 18–24 year old heterosexual Swiss adults who can be susceptible (*S*), infected (*I*), recovered (*R*) or vaccinated (*V*). These compartments are further divided into sub-compartments that reflect the individuals’ sex, sub-population/canton and sexual activity level, and can be described by the following system of ordinary differential equations (ODEs):

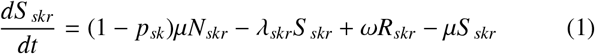

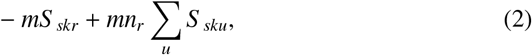

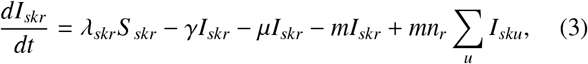

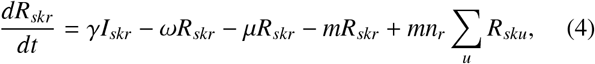

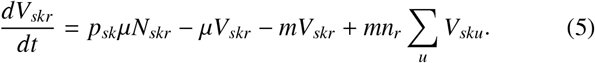

Here, the subscripts *s*, *k* and *r* denote sex, sub-population/canton and sexual activity group, respectively. Susceptible individuals (*S*) can become infected at rate *λ_skr_* (force of infection). Infected individuals (*I*) spontaneously clear HPV-16 at rate *γ* to become temporarily immune. Recovered individuals (R) loose their immunity at rate *ω* and become susceptible again. All individuals enter and leave the population at rate *μ* with *N_skr_* = *S _skr_* + *I_skr_* + *R_skr_* + *V_skr_* being the population size of individuals that have sex *s*, reside in sub-population/canton *k* and belong to sexual activity group *r*. *p_sk_* is the sub-population- or canton-specific proportion of individuals that are vaccinated upon entering the population. We assumed vaccine efficacy is 100% (3 doses) and lasts for an individual’s sexual lifetime. Individuals can change their sexual behavior at rate *m*, i.e., they are redistributed to either the same or another sexual activity group proportional to the size of the target group [19, 22].

### 2.2. Data and parameters

#### 2.2.1. Vaccination uptake

We used data from the Swiss National Vaccination Coverage Surveys (SNVCS) to obtain the proportion of women who are vaccinated in each canton (Fig. 1, Supplementary Material Table S.1). The SNVCS monitor immunization coverage of children and adolescents and compiles them into three-year bands. For HPV vaccination, the surveys focus on 16 years old girls. In this study, we used data from the last available survey period (2011–2013), except for the canton of Geneva (GE) and Appenzell Innerrhoden (AI) where we used data from the years 2010 and 2014, respectively. Two HPV vaccines are currently authorized in Switzerland: Gardasil^®^ (Sanofi Pasteur MSD) which targets four HPV types (HPV-6/11/16/18), and Cervarix^®^ (GlaxoSmithKline) which targets two HPV types (HPV-16/18). 95% of vaccinated women received the quadrivalent vaccine [4]. We used the proportion of fully vaccinated women (completed three doses) as a model parameter. Although Switzerland adopted the two-dose HPV vaccination schedule in 2012, we assumed that this had not been implemented in the cantonal programs at the time the surveys were done. We did not consider HPV vaccination in boys and young men, as uptake in Switzerland is negligible at present.

#### 2.2.2. Sexual behavior

We used data from the SIR (Screening, Impfung und Risikofaktoren) survey [4]. The Swiss Federal Office of Public Health (FOPH) conducted this survey in 2014 and collected data on the sexual behavior of 18–24 year old Swiss women (*n* = 1, 291). We categorised the study participants into two sexual activity groups and estimated the sexual partner change rates by assuming that the reported numbers of new heterosexual partners in the last year can be described by two Poisson distributions, weighted by the proportion of individuals in each sexual activity group [19, 23]. The survey did not include men, so we assumed their sexual activity to be the same as for women. Furthermore, we assumed that sexual behavior does not differ between cantons.

#### 2.2.3. Inter-cantonal mixing

We used mobility data from the Swiss Federal Office for Spatial Development (ARE) as a proxy for sexual mixing between different cantons. The data set contains average daily commuting data by public transport and individual vehicles from Monday to Friday in 2010 [24].

#### 2.2.4. Other parameters

We used publicly available data about the number of 18–24 year olds in each canton in 2013 from the website of the Swiss Federal Statistical Office (FSO) [25] (Supplementary Material Table S.2). Parameters that describe the transmission and life-history of HPV-16 were informed by the literature [26, 27] and assumed to be the same for women and men. All parameter values and their sources are specified in Table 1.

**Table 1.** Summary of parameters for the HPV-16 transmission model.

| Parameter | Description | Value | Unit | Reference/Comment |
| --- | --- | --- | --- | --- |
| $N_{skr}$ | Number of 18–24 year olds of sex $s$, sub-population/canton $k$ and activity group $r$ | See Table S.2 | – | Swiss FSO |
| $n_l$ | Proportion in the low sexual activity group | 0.85 | – | Estimated |
| $n_h$ | Proportion in the high sexual activity group | 0.15 | – | Estimated |
| $c_l$ | Heterosexual partner change rate in low activity group | 0.17 | per year | Estimated |
| $c_h$ | Heterosexual partner change rate in high activity group | 2.41 | per year | Estimated |
| $\mu$ | Rate at which individuals enter and leave the population | 0.14 | per year | 7-year age band |
| $m$ | Rate at which individuals can change activity groups | 1.0 | per year | [19, 22] |
| $\epsilon_r$ | Assortativity index for sexual mixing between activity groups | 0.5 | – | [19, 22] |
| $\epsilon_k$ | Assortativity index for sexual mixing between sub-populations/cantons | 0.6, 0.8, 1.0 | – | Assumption |
| $s$ | Scaling factor for mobility-informed sexual mixing matrix $\sigma_{kk'}$ | 0.035 | – | Calculated |
| $\beta$ | Transmission probability per partnership | 0.8 | – | [26] |
| $\gamma$ | Rate at which infection is cleared spontaneously | 0.55 | per year | [27] |
| $\omega$ | Rate at which immunity is lost | 0.024 | per year | [27] |
| $p_{sk}$ | Proportion of vaccinated individuals in canton $k$ | Fig. 1 | – | Swiss FOPH |

### 2.3. Sexual mixing and force of infection

The force of infection, *λ*_skr_, depends on assumptions about sexual contact preferences between individuals from different sexual activity groups and sub-populations/cantons. We devised three different scenarios of increasing complexity to account for different spatial mixing patterns (Fig. 2):

1. *Assortative sexual mixing:* Sexual contacts only occur between individuals from the same sub-population/canton.
2. *Proportional sexual mixing*: A fraction of sexual contacts occur between individuals from the same subpopulation/canton, while the remaining contacts are proportionally distributed across all sub-populations/cantons.
3. *Mobility-informed sexual mixing*: Swiss mobility data are used as a proxy for inter-cantonal sexual mixing.

**Fig. 2.**
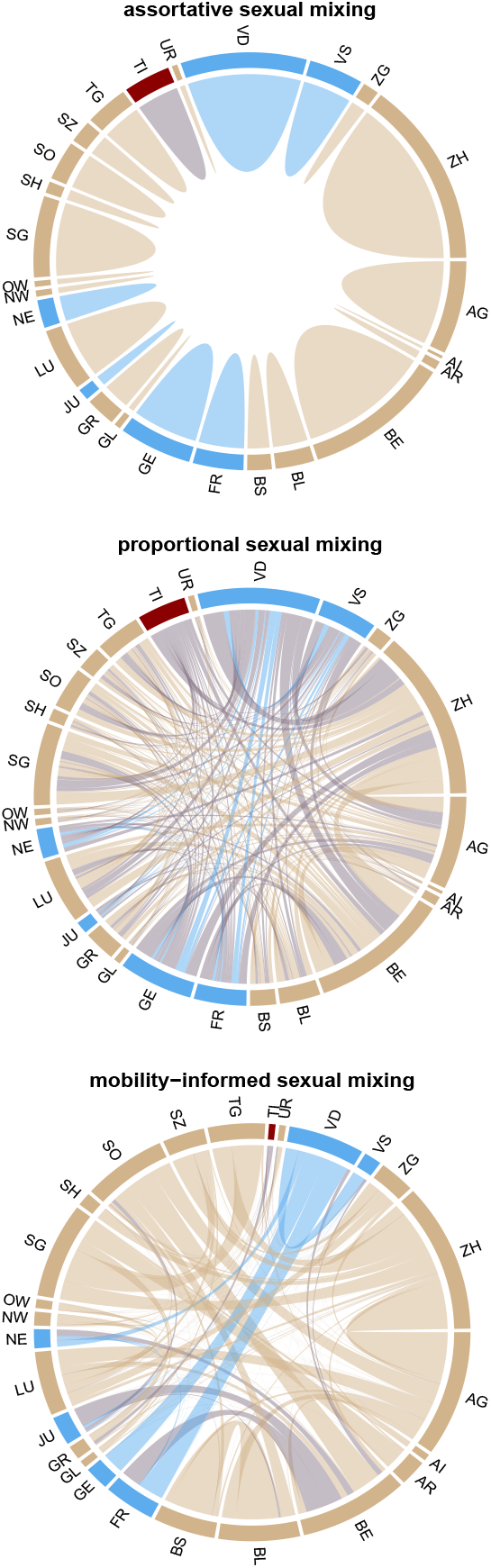
Chord diagrams of inter-cantonal sexual mixing. The diagrams show the number of sexual contacts between individuals from different cantons. For the scenarios where sexual mixing between cantons occurs (proportional and mobility-informed sexual mixing), we excluded the sexual contacts between individuals that reside in the same canton for better visibility. Cantons with a French-, German- or Italian-speaking majority are indicated in blue, beige and red, respectively. Acronyms for canton names are explained in the Supplementary Material Table S.1

#### 2.3.1. Assortative and proportional sexual mixing

The first two scenarios where we assumed fully assortative or partial proportional mixing between sub-populations/cantons result in the following force of infection:

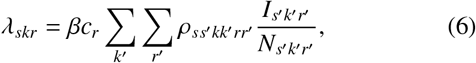

where *β* is the per partnership transmission probability and *c_r_* is the sexual partner change rate for individuals of sexual activity group *r*. The elements of the sexual mixing matrix

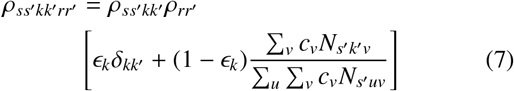

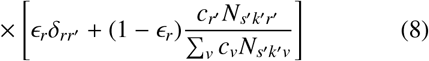

describe the conditional probability of an individual of sex *s*, sub-population/canton *k* and sexual activity group *r* to have a sexual contact with an individual of the opposite sex *s*′, sub-population/canton *k*′ and sexual activity group *r*′. *∈_k_* and *∈_r_* are the sexual mixing coefficients with respect to sub-population/canton and sexual activity group, respectively. Values of 1 represent fully assortative mixing where individuals only have sexual contacts with other individuals from the same sub-population/canton or sexual activity group. A value of 0 corresponds to proportional (random) mixing where sexual partners are chosen in proportion to the size of their sub-population/canton and their sexual activity group. *δ_kk′_* and *δ_rr′_* are the Kronecker deltas that are equal to 1 if *k* = *k*′ or *r* = *r*′ and to 0 otherwise. In the first scenario (assortative sexual mixing), we set *∈_k_* = 1. In the second scenario (proportional sexual mixing), we set *∈_k_* to 0.6 (model with two sub-populations) and 0.8 (cantonal model). Throughout all simulations, we set *∈_r_* = 0.5, which corresponds to partially assortative mixing with respect to sexual activity [19, 22].

#### 2.3.2. Mobility-informed sexual mixing

We used mobility data as a proxy for inter-cantonal sexual mixing by assuming that the heterosexual partner preference across cantons is proportional to the corresponding commuting patterns. The symmetrical matrix *P*_mob_ provides absolute numbers of commuters between cantons without specifying the commuters’ canton of residence. We converted *P*_mob_ into an asymmetrical inter-cantonal mixing matrix *σ_kk′_* that provides the conditional probabilities that a sexual contact from an individual from canton *k* occurs with someone from canton *k*′. To this end, we first rescaled *P*_mob_ by a scaling factor *s* and weighted all columns with the inverse of the cantonal population size:

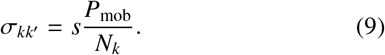

We then replaced the diagonal entries of *σ_kk′_* with the sum of, all entries that are outside canton *k*:

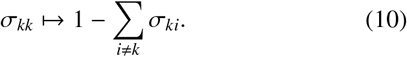

The force of infection for the mobility-informed sexual mixing scenario is given by Eq. 6 with *ρ_ss′kk′rr′_* being replaced by *σ_kk′_ρ_ss′rr′_*. We chose the scaling factor *s* such that the weighted proportion of intra-cantonal heterosexual contacts across all cantons is 80% (Supplementary Material Fig. S.1), i.e., is the same as in the proportional sexual mixing scenario:

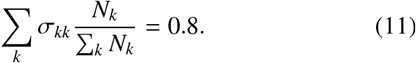

### 2.4. Model simulations

We simulated the different model scenarios by numerically integrating the ODEs until the system approached the endemic pre-vaccination equilibrium (*p_sk_* = 0). We then initiated the HPV vaccination program by setting *p_sk_* > 0, and ran the model for a further number of years. The ODEs were solved in the R software environment for statistical computing [28] using the function *ode* from the package *deSolve*. We calculated the basic reproduction number (*R*_0_) using the next-generation matrix method as described by Diekmann et al. [29, 30] (Supplementary Material Section 1). This allowed us to compute the vaccination threshold *V_C_* = 1 − 1/*R*_0_. All code files can be downloaded from GitHub

## 3. Results

### 3.1. HPV-16 dynamics

Using the parameters from Table 1, the transmission model provides a realistic description of the HPV-16 dynamics in Switzerland. The pre-vaccination prevalence of HPV-16 is 3.34% among 18–24 year olds. While this is somewhat lower than the expected and observed HPV-16 prevalence in Britain (Supplementary Material Section 2, Table S.3), it is in the range that is typically observed among women in other European countries [20]. The functional relationship between vaccination coverage and the reduction in HPV-16 prevalence 2 to 4 years post-vaccination is in good agreement with the findings of a systematic review (Supplementary Material Section 3, Fig. S.2) [1]. The basic reproduction number, *R*_0_, of HPV-16 in our model is 1.29. This value corresponds to a vaccination threshold of 22% in the general population. If vaccination is targeting only one sex, the threshold increases to 39%.

### 3.2. Vaccination in two sub-populations

To better understand the effects of spatially heterogeneous vaccination uptake on infection transmission, we focused on a simplified model with just two sub-populations of the same size. We calculated the expected HPV-16 prevalence after 50 years of vaccinating the two sub-populations at different coverage rates (Fig. 3). In the first scenario, we assumed fully assortative sexual mixing between the two sub-populations, i.e., sexual contacts only occur between individuals from the same sub-population (Fig. 3a). The concave relation between vaccination coverage in the two sub-populations and the expected prevalence of HPV-16 overall indicates that homogeneous vac-; cination uptake always has the largest effect on reducing prevalence. For example, a vaccination coverage of 25% in both sub-populations results in a lower prevalence than vaccinating either of them at 50%. In the second scenario, we assumed a certain level of proportional mixing where 20% of sexual contacts are made with individuals from the other sub-population (Fig. 3b). Sexual mixing between the two sub-populations diminishes the negative effect of heterogeneous vaccination uptake, but homogeneous vaccination still results in the lowest prevalence of HPV-16. Fig. 3c shows the difference in the expected HPV-16 prevalence between the first (no sexual mixing; between the sub-populations) and second (sexual mixing between the sub-populations) scenario. The higher the difference, the stronger the effect of sexual mixing is in reducing the negative consequences of heterogeneous vaccination uptake. This is particularly the case when vaccination is highly heterogeneous, i.e., when uptake is very high in one sub-population and very-low in the other sub-population. In summary, these results illustrate that spatially heterogeneous vaccination uptake diminishes the effect of vaccination on reducing HPV-16 prevalence, but that sexual mixing between sub-populations can limit these undesired consequences by ‘homogenizing’ the overall population.

**Fig. 3.**
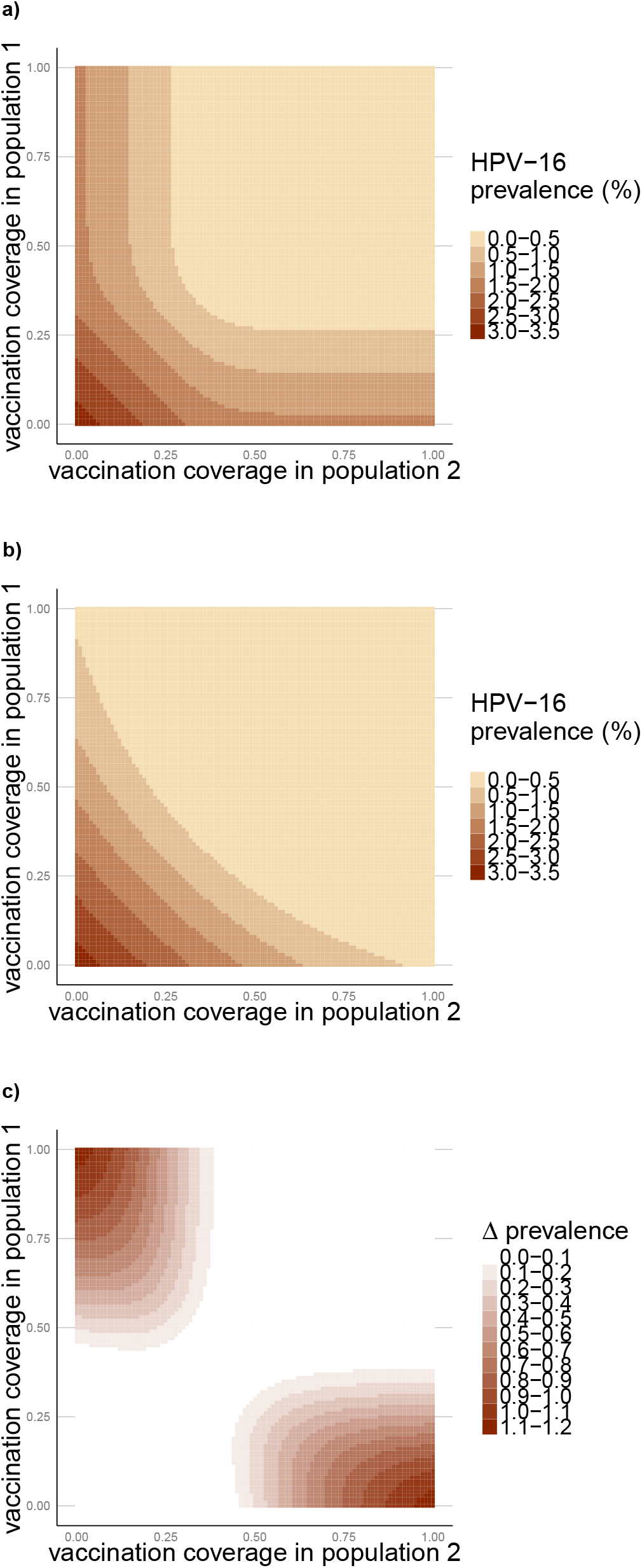
Heterogeneous vaccination uptake and HPV-16 prevalence. The graphs show the expected prevalence of HPV-16 after 50 years of vaccinating two subpopulations at different coverage rates. a) HPV-16 prevalence when there is no sexual mixing between the two populations. b) HPV-16 prevalence when 20% of sexual contacts are made between the two populations (*ε_k_* = 0.6). c) Difference in HPV-16 prevalence between scenario a and b.

### 3.3. Transmission of HPV-16 within and between cantons

We extended our analysis of heterogeneous vaccination uptake by simulating the transmission of HPV-16 within and between the 26 cantons of Switzerland. The observed dynamics generalize some of the insights from the simplified model with two-subpopulations. After vaccination is introduced, HPV-16 prevalence begins to diverge across cantons (Fig. 4). After 15 years of vaccination, the range of expected HPV-16 prevalences depends on the assumed scenario for sexual mixing between cantons (see Methods). For fully assortative mixing, the highest and lowest prevalence are 2.40% (ZG, 17% vaccination coverage) and 0.12% (VS, 75% vaccination coverage), respectively (Fig. 4a). The range of cantonal HPV-16 prevalence narrows if sexual mixing between cantons is taken into account. The cantonal prevalence ranges from 1.28% to 0.23% for proportional mixing (Fig. 4b), and from 1.09% to 0.14% for mobility informed mixing (Fig. 4c). Thus, sexual mixing between cantons again ‘homogenizes’ the infection dynamics and the effect of vaccination on reducing prevalence.

**Fig. 4.**
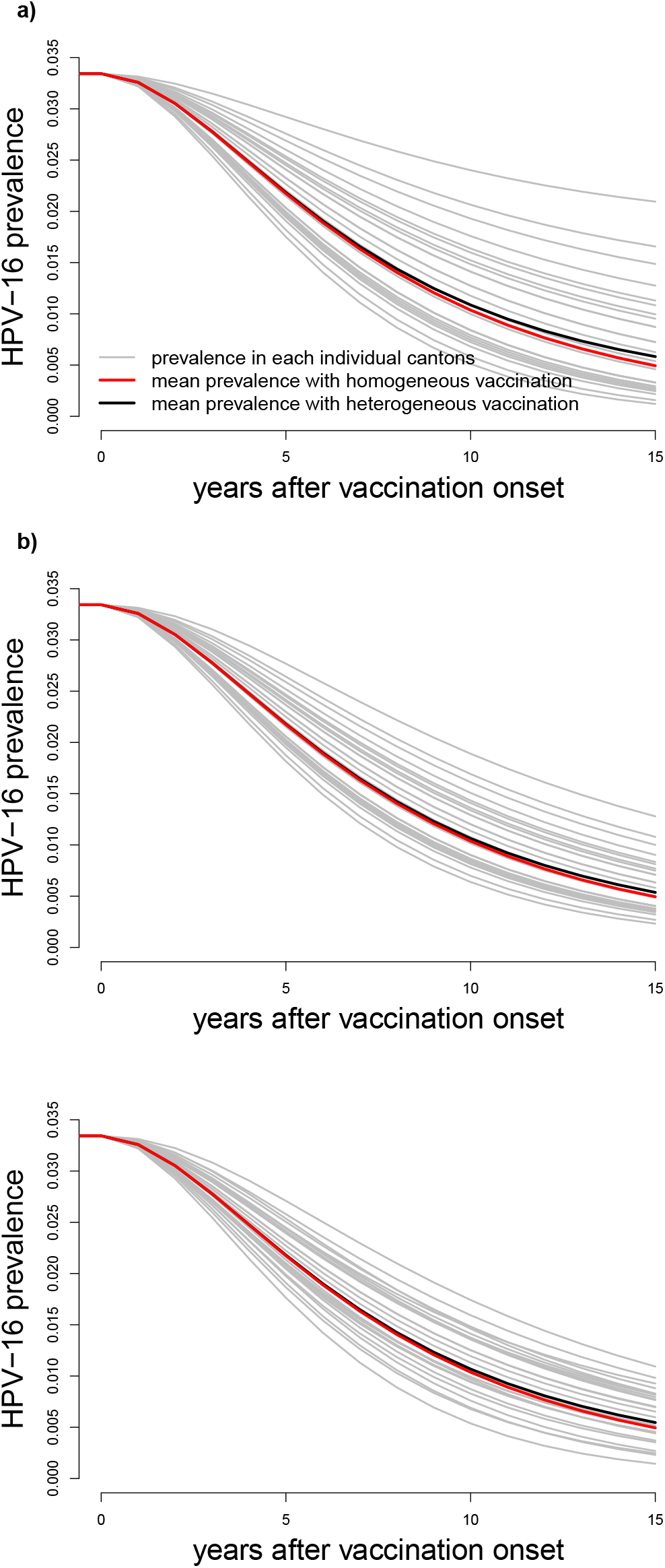
Cantonal and national prevalence of HPV-16 after vaccine introduction. a) Fully assortative mixing (no sexual mixing between cantons). b) Proportional mixing (20% of sexual contacts are proportionally distributed over all of Switzerland). c) Mobility-informed mixing. Grey lines represent cantonal HPV-16 prevalence. The black and red lines correspond to the national prevalence for heterogeneous and homogeneous vaccination uptake, respectively.

This effect is also reflected in the overall prevalence of HPV-16 in Switzerland. The national prevalence of HPV-16 is slightly higher under heterogeneous vaccination uptake compared with homogeneous uptake (Fig. 4a). This difference becomes smaller in the two scenarios that assume sexual mixing between the two cantons (Fig. 4b and 4c). In the most realistic scenario (mobility-informed mixing), the national prevalence of HPV-16 is expected to drop to 0.55% after 15 years of heterogeneous vaccination uptake, while homogeneous vaccination uptake would drop the prevalence to 0.49%. The result that heterogenous vaccination uptake yields a slightly higher HPV-16 prevalence compared with homogeneous uptake is robust to different assumptions about sexual activity, cantonal population sizes and the overall vaccination uptake ((Supplementary Material, Table S.4)).

Inter-cantonal sexual mixing helps to reduce the prevalence of HPV-16 in cantons with low vaccination coverage at the expense of cantons with high vaccination coverage. At the national level, increasing sexual mixing between cantons always results in a lower HPV-16 prevalence (Fig. 5, dashed red lines), while the effect of sexual mixing at the cantonal level is more intricate. The number of cantons that achieve a specific reduction in prevalence – expressed as relative risk (RR) reduction – can either decrease or increase with varying degrees of sexual mixing (Fig. 5). For example, high levels of sexual mixing between cantons (low *∈_k_*) increase the number of cantons that achieve a 50% reduction in prevalence after 15 years of vaccination (Fig. 5a). In contrast, low levels of sexual mixing between cantons (high *∈_k_*) are required to increase the number of cantons that achieve a RR reduction of 90%. On a timescale of 50 years, the number of cantons that reach a RR reduction of 99% is lowest for low, but realistic, levels of sexual mixing between cantons (*∈_k_* = 0.85 − 0.95) (Fig. 5b). These levels of sexual mixing prevent the elimination of HPV-16 in high-coverage cantons, but they are too low for low-coverage cantons to sufficiently benefit from the herd immunity of high-coverage cantons.

**Fig. 5.**
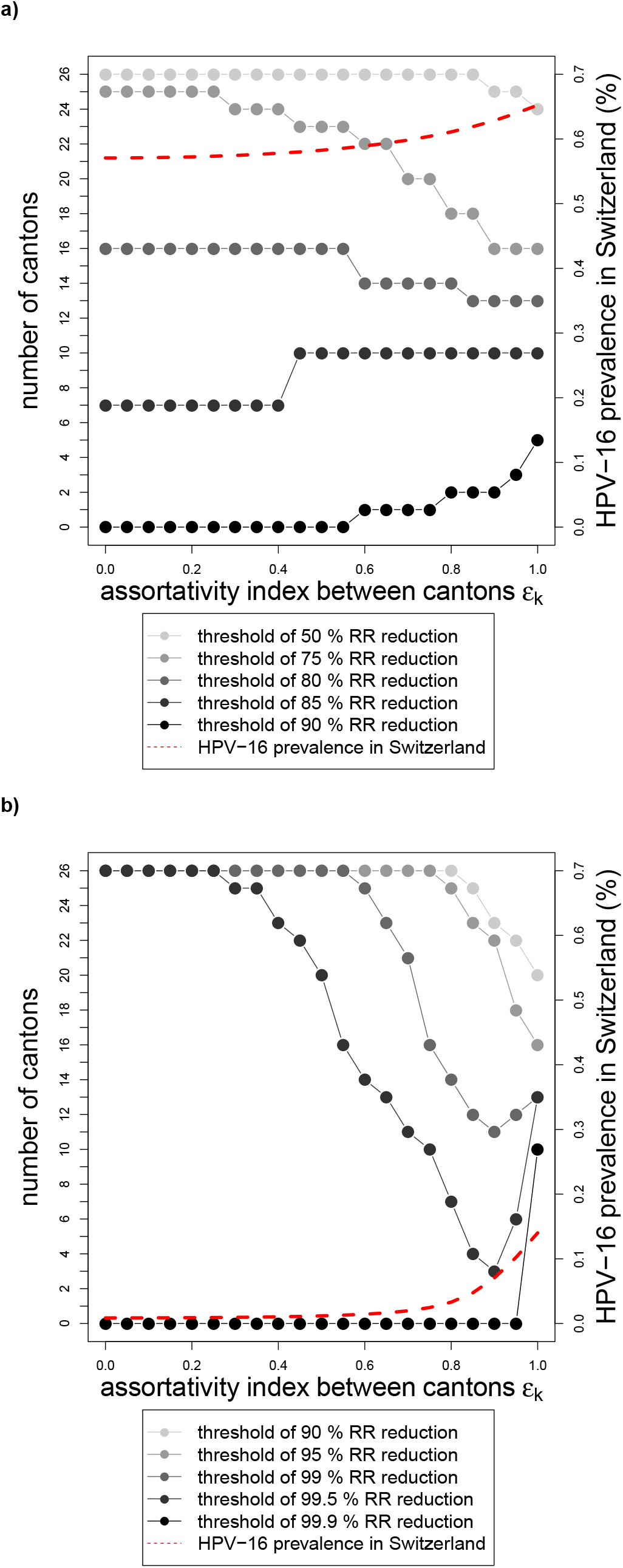
Relationship between inter-cantonal sexual mixing and HPV-16 prevalence. The graphs show the number of cantons that achieve a specific relative risk (RR) reduction after 15 years (a) and 50 years (b) of vaccination. The dashed red lines correspond to the national prevalence which is lowest if sexual mixing is completely proportional (*∈_k_* = 0). For all simulations, we used the proportional sexual mixing scenario.

## 4. Discussion

Uptake of HPV vaccination in 16 year old girls in Switzerland shows pronounced differences between different cantons ranging from 17 to 75%. We used a dynamic transmission model to study the expected consequences of this spatial heterogeneity in vaccination uptake on the transmission and prevalence of HPV-16 in Switzerland. Using a simple model with just two sub-populations, we found that heterogeneous vaccination uptake can diminish the effect of vaccination on reducing HPV-16 prevalence. This effect is strongest when vaccination is highly heterogeneous, i.e., when uptake is very high in one sub-population and very low in the other sub-population. These results were then corroborated with an extended model simulating the transmission of HPV-16 within and between the 26 cantons of Switzerland. Homogeneous vaccination uptake would generate a lower national HPV-16 prevalence compared to heterogeneous vaccination uptake, but the overall differences in prevalence are very small. We found that inter-cantonal sexual mixing ‘homogenizes’ the infection dynamics, limits the undesired consequences of heterogeneous vaccination uptake, and reduces the inter-cantonal differences in HPV-16 prevalence.

This study describes the transmission of HPV-16 in Switzerland using a mathematical model to investigate how spatial heterogeneity in vaccination uptake affects prevalence. The example of Switzerland provides sufficient data for parameterizing a dynamic transmission model while exhibiting large variation in HPV-16 vaccine deployment. Using Swiss sexual behavior data, the model provided a realistic description of HPV-16 transmission in Switzerland, and allowed us to investigate the expected effect of HPV vaccination. Our results do not change qualitatively when the number of cantons or parameter values are varied within reasonable ranges. In the absence of data describing inter-cantonal sexual mixing in Switzerland, we used commuting data and explored three different scenarios. The two scenarios that assumed partial sexual mixing between cantons – proportional sexual mixing and mobility-informed mixing – gave rise to a similar pattern, strengthening the validity of our findings.

Our study has a number of limitations that need be considered when interpreting the findings. First, we used a relatively simple model to describe the transmission of HPV-16, not taking into account potential sex-specific differences in sexual behavior and the infection life-history. Owing to our focus on the transmission and prevalence of HPV-16, we did not include the progression of HPV infections to cervical intraepithelial neoplasia (CIN), as some modeling studies have done [7, 8, 26, 27]. Further, we did not consider different age classes and assumed that women can only become vaccinated before the age of 18. It is also important to note that our results depend on the assumption that the sexual behavior and the subsequent risk of HPV infection is the same across different cantons. Second, comparing HPV-16 prevalence and the sexual behavior data (i.e., the estimated heterosexual partner change rates) between Swiss and British women needs to be treated with caution. Although the particular question about the number of new heterosexual partners was the same in both surveys, the methods for sampling and data collection differed considerably. While the SIR survey interviewed participants by phone, Natsal-3 relied on individuals filling in questionnaires at the participants’ homes. This difference could have introduced a social desirability bias that could result in an underestimation of heterosexual partner changes based on the SIR study. This underestimation might be further compounded by the fact that the SIR survey included only women. Given the sensitivity of our model with regard to per partnership transmission probabilities (Supplementary Material Fig. S.3) and heterosexual partner change rates, our calculations of *R*_0_ and the corresponding vaccination thresholds should therefore be interpreted with caution. Third, in absence of data about the levels of sexual mixing between cantons, we assumed that inter-cantonal sexual mixing is proportional to the observed commuting patterns. Furthermore, we assumed that the national average of sexual contacts that are made with individuals from the same canton is 80%, and that 20% are made with individuals from another canton. This assumption was informed by a Canadian study on couple composition regarding language membership (French, English or other) led in 1981 [31]. On average, 18.2% of couples in Quebec were exogamous, with some heterogeneity over different regions. Fourth, besides inter-cantonal variation in HPV vaccination uptake, there is also intra-cantonal variation. For example, vaccination uptake in Geneva, which has a school-based vaccination program, varies significantly among different nationalities and socio-economical status [32]. Investigating the causes and consequences of intra-cantonal variation in HPV vaccination uptake in Switzerland is part of ongoing work.

There are currently no population-based prevalence estimates of type-specific HPV in Switzerland. Our modeled prevaccination prevalence of HPV-16 is 3.34%, and is within a plausible range for women in European countries. A metaanalysis of more than 1 million women estimated HPV-16 prevalence at 4.8% and 3.2% in Europe and globally, respectively [20]. Only a few studies provide estimates for the basic reproduction number, *R*_0_, or equivalently, the vaccination threshold of HPV-16 or other HPV types. Ribassin-Majed et al. [33] estimated *R*_0_ = 1.73 for HPV-16/18 in France, corresponding to a vaccination threshold of 67% for one sex. These values are higher than what we calculated for Switzerland, but in a similar range to what would be expected in Britain (Table ??). The lower values that we calculated for Switzerland underline the possibility of underreporting in the Swiss sexual behavior survey.

Our results need to be interpreted in the context of the current HPV literature considering heterogeneity in vaccination. The finding that decreasing heterogeneity in vaccination uptake increases impact helps to interpret the result by Durham et al. [15] who showed that vaccination efforts should be targeted towards low-vaccination states in the USA. Increasing vaccination uptake in populations with low-vaccination uptake has the strongest effect for reducing vaccination heterogeneity overall. The study by Shafer et al. [16] on unequal HPV vaccination uptake among different ethnic groups in Canada, suggests that heterogeneous vaccination can lead to cross-over effects across groups and depends on the amount of sexual mixing between the groups. Our study corroborates these findings and illustrates the effect of heterogeneous vaccination uptake between different populations and its relationship with different amount of sexual mixing between them.

We showed that the effect of cantonal variations in vaccination uptake on reducing the overall effect of vaccination on HPV-16 prevalence in Switzerland is small. This result is remarkable as eight cantons (ZG, AR, SZ, OW, AI, TG, BE and TI) have a vaccination uptake that is below the vaccination threshold in our model (39.6%). In contrast, all cantons are above this threshold assuming homogeneous vaccination at an uptake that is identical to the national average (52%). One might expect that the effects of herd immunity in the latter scenario would result in a substantially lower prevalence of HPV-16 compared with heterogenous uptake. However, we compared the expected prevalence after 15 years of vaccination when prevalence is still declining rapidly in all cantons and the post-vaccination equilibrium has not been reached. The rapid and pronounced decline in HPV-16 prevalence that we found in our model even for populations with low vaccination uptake is in good agreement by epidemiological studies [1, 34].

Our findings could have implications for the future planning of HPV vaccination programs at the cantonal and national level in Switzerland. From the point of view of a particular canton, the achieved reduction in HPV-16 prevalence will not only depend on the cantonal vaccination program, but also on the indirect effects of vaccination efforts in other (particularly neighboring) cantons and how these effects are dissipated via intra-cantonal sexual mixing. For the most plausible scenario for inter-cantonal mixing (mobility-informed sexual mixing), we found that cantons with high vaccination coverage experience a less effective reduction in HPV-16 prevalence to what would be expected if they were isolated (assortative sexual mixing). Conversely, this effect benefits those cantons with a low vaccination uptake that achieve a higher reduction in prevalence to what would be expected in absence of intra-cantonal sexual mixing. The intensity of cantonal dissipation of vaccination efforts is again mediated by intra-cantonal sexual mixing. The number of cantons that surpass a pre-defined relative risk reduction is highly sensitive to the level of assortative mixing between cantons (Fig. 5). The results of this study suggest that a harmonization of programs between cantons, and a reduction in vaccination heterogeneity, would result in a stronger effect of vaccination on reducing HPV-16 prevalence in Switzerland. The generality of our results on the effects of spatial heterogeneity in vaccination uptake are also relevant for the planning of vaccination programs in other countries, and in the context of infectious diseases other than HPV.

In summary, we found that spatial heterogeneity in HPV vaccination uptake is expected to diminish the effect of vaccination on HPV-16 prevalence, but the overall effect is small. In the context of Switzerland, this means that cantonal efforts towards a reduction of HPV-prevalence are impaired by cantons with low vaccination uptake. Harmonization of cantonal vaccination programs would reduce inter-cantonal differences in HPV-16 prevalence.

## Acknowledgment

We would like to thank the Swiss Federal Office of Public Health (FOPH), the Swiss Federal Office for Spatial Development (ARE), and the investigators of the British National Survey of Sexual Attitudes and Lifestyles (Natsal) for providing. access to the data used in this study. We would also like to thank J.A. Bogaards for his valuable comments on our study.

